# The lncRNA SOX2OT Drives Non-Small Cell Lung Cancer Progression and Metastasis by Suppressing miR-143

**DOI:** 10.64898/2026.07.27.741140

**Authors:** Mohadeseh Zarei, Elahe Asadollahi, Babak Jahangiri, Jamshid Raheb

**Affiliations:** Department of Molecular Medicine, National Institute of Genetic Engineering and Biotechnology, Tehran, Iran; Department of Genetics, Faculty of Biological Sciences, Tarbiat Modares University, Tehran, Iran

**Keywords:** SOX2OT, miR-143, non-small cell lung cancer, lncRNA, epithelial-mesenchymal transition, cancer progression

## Abstract

In terms of cancer-related death, non-small cell lung cancer (NSCLC), the world’s leading cause, highlights the need for continued research into the genetic factors that influence tumor growth. Long non-coding RNAs (lncRNAs) are now well recognized as essential regulators of oncogenic signaling cascades; nevertheless, the specific role and molecular basis of the SOX2 overlapping transcript (SOX2OT) in NSCLC are not entirely understood. This study examined the functional importance of SOX2OT and its regulatory interactions with tumor-suppressive microRNAs in NSCLC cells. In A549 and Calu-3 cells, RNA interference-mediated SOX2OT silencing dramatically reduced cellular proliferation, migration, and invasiveness. Moreover, SOX2OT knockdown was associated with inhibition of epithelial–mesenchymal transition (EMT), alongside induction of cell cycle arrest and activation of apoptotic pathways. Integrated transcriptomic profiling and bioinformatic prediction analyses identified miR-143 as a putative downstream effector of SOX2OT activity. Consistently, depletion of SOX2OT resulted in marked elevation of miR-143 expression, which corresponded with downregulation of oncogenic mediators, including STAT3, EZH2, and CXCL13. As a result of SOX2OT suppression, both the transcript and the protein levels of PTEN were restored. Further functional characterization demonstrated that SOX2OT knockdown inhibits EMT progression by decreasing mesenchymal markers and EMT-related transcription factors (TFs) while concomitantly enhancing epithelial marker expression. Collectively, these findings suggest that SOX2OT contributes to NSCLC pathogenesis through regulation of a miR-143-centered signaling network that influences oncogenic signaling, cellular survival, and metastatic potential. Targeting the SOX2OT/miR-143 regulatory axis may therefore represent a promising therapeutic approach for NSCLC, while also underscoring the broader importance of lncRNA-mediated post-transcriptional regulation in lung cancer biology.

## Introduction

Despite significant progress in targeted therapies and immunotherapeutic strategies, non-small cell lung cancer (NSCLC) continues to represent a major global health challenge. Lung cancer accounts for nearly 85% of all cancer diagnoses worldwide, making it the leading cause of cancer-related death (1–3). The continually poor prognosis of NSCLC patients, particularly those with advanced or metastatic cancer, underscores the critical need to understand the molecular pathways that promote tumor development and therapy resistance (4). A deeper understanding of the regulatory networks governing NSCLC malignancy may enable the identification of novel molecular vulnerabilities and improve clinical outcomes.

Long non-coding RNAs (lncRNAs) have emerged as critical regulators of gene expression at multiple levels — transcriptional, post-transcriptional, and epigenetic — playing pivotal roles in cancer development and progression (5). Through interactions with nucleic acids and proteins, lncRNAs modulate key biological processes including proliferation, apoptosis, migration, invasion, and EMT. Given the growing link between lncRNA dysregulation and NSCLC pathogenesis, these molecules are increasingly recognized as valuable biomarkers and targets for therapy (6).

SOX2 overlapping transcript (SOX2OT) is a well-characterized oncogenic lncRNA transcribed from the SOX2 locus, a genomic region critical for stemness and cellular plasticity (7, 8). The aberrant expression of SOX2OT has been documented across various malignancies, particularly in lung cancer. In this context, SOX2OT is strongly implicated in driving tumor growth, invasion, resistance to therapy, and immune evasion (9). SOX2OT exerts regulatory control over cancer-associated signaling pathways through various mechanisms, including the modulation of SOX2 expression and interactions with microRNAs (miRNAs) (10–12). However, the molecular pathways by which SOX2OT contributes to NSCLC progression remain incompletely defined.

MiRNAs are small non-coding RNAs that post-transcriptionally regulate gene expression by targeting complementary sequences in messenger RNAs, thereby influencing cellular fate decisions (13). Among them, miR-143 has consistently been reported as a tumor-suppressive miRNA across multiple cancer types, including lung cancer (14, 15). NSCLC cells with reduced miR-143 expression are more likely to proliferate, migrate, invade, undergo apoptosis resistance, and migrate during disease progression(16, 17). Restoration of miR-143 expression potently suppresses malignant phenotypes in NSCLC cells by directly targeting multiple oncogenic regulators, underscoring its critical role as a tumor suppressor (16).

Emerging evidence suggests that lncRNAs may function as competing endogenous RNAs (ceRNAs) by sequestering miRNAs and thereby relieving repression of oncogenic targets (18). Increasing evidence suggests that SOX2OT plays a role in the progression of various cancers by functioning as a competing endogenous RNA (ceRNA), effectively trapping tumor-suppressive microRNAs such as miR-122-3p, miR-194-5p, and miR-30d-5p. This interaction leads to the derepression of oncogenic signaling cascades, thereby driving malignant progression (9). Notably, recent studies have suggested a potential interaction between SOX2OT and miR-143 in hepatocellular carcinoma, implicating this regulatory axis in tumor progression (19). Whether a similar SOX2OT/miR-143-mediated mechanism operates in NSCLC remains unexplored.

In this study, we attempted to clarify SOX2OT’s functional involvement in NSCLC and its possible interaction with miR-143 and downstream oncogenic signaling pathways. We used the A549 and Calu-3 NSCLC cell lines to investigate the effects of SOX2OT silencing on cellular proliferation, cell cycle progression, apoptosis, migration, invasion, and EMT. Additionally, we investigated the effects of SOX2OT knockdown on miR-143 levels and its downstream targets, specifically STAT3, EZH2, CXCL13, and PTEN. These results elucidate a novel SOX2OT-mediated regulatory network in NSCLC, underscoring the therapeutic promise of targeting the SOX2OT/miR-143 axis in lung cancer management.

## Material and methods

### Ethics statement

Throughout the investigation, all guidelines and regulations applicable to institutional and international research were followed. We sourced Calu-3 and A549 human NSCLC cell lines from the Pasteur Institute (Tehran, Iran). These immortalized cell lines were originally established from patient-derived material obtained following informed consent. Iranian National Institute of Genetic Engineering and Biotechnology’s Institutional Ethics Committee evaluated and approved all experimental protocols. No primary human tissues or directly patient-derived specimens were utilized in this study.

### Cell culture

The NSCLC cell lines Calu-3 and A549 were grown in DMEM (Gibco) supplemented with 10% fetal bovine serum (FBS) and 1% penicillin-streptomycin. Cultures were incubated at 37°C in a humid environment with 5% CO₂. Cell viability and morphology were consistently examined at stages 4-10.

### siRNA transfection

GenScript provided SOX2OT-targeting small interfering RNAs (si-SOX2OT) and a non-targeting scrambled siRNA to serve as a negative control. To assess transfection effectiveness, the control siRNA was coupled with 5’-fluorescein (6-FAM). 3 × 10⁵ cells were seeded in each well of 6-well culture plates and cultured overnight to permit adequate cell adherence. Transfections were performed at about 70% confluence using Lipofectamine 2000 agent (Invitrogen) according to the manufacturer’s instructions. Before transfection (2 hours), the culture media was changed with decreased serum medium containing 5% FBS and antibiotics. All experimental conditions were carried out in triplicate.

### RNA extraction and qRT-PCR analysis

Calu-3 and A549 cells (untreated and transfected) were isolated with RNX-Plus reagent (Sinaclon, Iran) 48 hours post-transfection. A NanoDrop spectrophotometer (Thermo Fisher Scientific) was used to quantify and purify RNA, while agarose gel electrophoresis was used to verify integrity. Contaminating genomic DNA was eliminated by DNase I treatment (Thermo Fisher Scientific). Using a mix of oligo(dT) and random hexamer primers, 2 μg of total RNA was reverse-transcribed into complementary DNA (cDNA) for mRNA and lncRNA studies using the AddScript cDNA Synthesis Kit (AddBio, Korea). BioFACTTM 2× Real-Time PCR Master Mix (BIOFACT, South Korea) was then used on a Mic qPCR Cycler system to perform quantitative real-time PCR. The expression quantities of SOX2OT and principal EMT markers (E-cadherin, N-cadherin, Vimentin, SNAIL, and TWIST) were normalized using GAPDH as the internal control gene. Stem-loop reverse transcription primers were used to create miR-143-specific cDNA for microRNA analysis. With SNORD48 acting as the endogenous control, SYBR Green-based qRT-PCR was used to quantify miR-143 under ideal cycling conditions. Every reaction was carried out three times. The 2^⁻ΔΔCt^ technique was used to compute relative gene expression, which was then shown as fold changes in comparison to control samples.

### Western blot analysis

The cells were lysed with RIPA buffer supplemented with a protease inhibitor mixture (Roche). Polyvinylidene difluoride (PVDF) membranes were used to separate equal amounts of total protein using SDS-PAGE. Following blocking membranes by 3% bovine serum albumin (BSA) for an hour at room temperature, primary antibodies against STAT3, PTEN, and CXCL13 (Santa Cruz Biotechnology; 1:1000 dilution) were incubated overnight at 4°C (8). Following washing, membranes were incubated for one hour at room temperature with a secondary antibody conjugated with horseradish peroxidase (HRP) (Santa Cruz Biotechnology, sc-2357; 1:5000 dilution). Immunoreactive proteins were detected using a chemiluminescence substrate (Thermo Fisher Scientific) and quantified using ImageJ software. As an internal loading control, β-actin was used to standardize relative protein expression levels.

### Cell cycle analysis

To evaluate cell cycle distribution, 48 hours after transfection the collected cells were preserved in 70% chilled ethanol and kept overnight at 4°C. PI/Triton X-100 staining solution (10 g/mL propidium iodide (PI), 100 g/mL RNase A, 0.1% Triton X-100) was used after washing with Phosphate-buffered saline (PBS). Samples were analyzed on a BD FACSCalibur flow cytometer (BD Biosciences), and data interpretation was performed using FlowJo software.

### Apoptosis assays

The apoptotic cell death was measured through dual staining with Acridine orange/ethidium bromide (AO/EtBr) and flow cytometry with Annexin V-FITC/PI. A549 and Calu-3 cells transfected with si-SOX2OT were compared to their respective control groups 48 hours after transfection. The cells were rinsed twice with PBS before they were resuspended in PBS for AO/EtBr labeling. A fresh staining solution comprising 100 μg/mL of EtBr and 100 μg/mL of AO in a 1:1 ratio had been added to the cell suspension. Fluorescence microscopy was used to see the cells right away. Early apoptotic cells had vivid green nuclear staining, late apoptotic or necrotic cells had orange to red fluorescence, and viable cells had uniform green fluorescence in the nucleus. A 200× magnification was used to get representative photographs. Using an Annexin V-FITC/PI apoptosis detection kit (BD Biosciences) and the manufacturer’s instructions, a quantitative assessment of apoptosis was carried out. In short, cells were extracted 48 hours after transfection, cleaned with PBS, and stained with Annexin V-FITC and PI for 15 minutes at room temperature in the dark. Flow cytometry was used to assess the samples right away, and FlowJo software was used to quantify the early and late phases of apoptotic populations. Every experiment was carried out three times.

### Cell proliferation and viability assays

Proliferation and viability were assessed via MTT and trypan blue assays. In the MTT assay, A549 and Calu-3 cells (9×10^3^ cells/well in 96-well plates) were transfected with si-SOX2OT or scrambled siRNA after overnight adhesion. Following 24 and 48 hours of incubation, MTT (2 mg/mL) was introduced for 4 hours at 37°C. Absorbance at 570 nm was measured after solubilizing formazan crystals in DMSO. Trypan blue exclusion was achieved by transfecting 7×104 cells in 24-well plates for 48 hours, harvesting and staining the cells, and manually counting the live cells. All test was run in triplicate.

### Cell migration assays

Cell migration potential was evaluated using both Transwell migration and wound-healing (scratch) assays. For the scratch assay, Calu-3 and A549 cells were plated into 6-well plates and maintained until reaching nearly 90% confluence. A uniform linear scratch was then created through the monolayer with a sterile 200-µL pipette tip. Following two rinses with PBS to eliminate cell debris, the cells were maintained in a culture environment supplemented with 1–2% FBS to minimize cell proliferation. Under 100× magnification, pictures were taken at 0, 24, and 48 hours. Using ImageJ software, the extent of wound healing was quantified using the following formula: [(original wound area − wound area at the selected time point) / initial wound area] × 100. Every experiment was carried out in triplicate, and each wound region had at least three randomly chosen areas examined.

For the transwell migration experiment, cells were collected 48 hours after transfection and plated into the upper chambers of Transwell inserts (8.0-µm pore diameter) at a density of 5 × 10² cells in 250 µL of serum-free DMEM. The lower chambers were loaded with 750 µL of complete medium comprising 10% FBS to serve as a chemoattractant. After a 48-hour incubation at 37°C, non-migrated cells left on the upper surface of the membrane were gently cleared using a cotton swab. Cells that had traversed to the underside of the membrane were subsequently fixed with 3.7% paraformaldehyde for 20 minutes, stained with Giemsa for 15 minutes, and quantified by counting six randomly selected microscopic fields per insert at 200× magnification.

### Cell invasion assay

A Matrigel-coated transwell test was used to measure invasion. Matrigel (BD Biosciences) was used to pre-coat inserts (8.0-µm pore size), which were then polymerized at 37°C. resuspended in a serum-deprived solution were placed into the top wells, whereas the bottom wells were supplemented with complete medium containing 10% FBS. Non-invading cells were eliminated from the top surface after a 48-hour incubation period. After fixing with 3.7% paraformaldehyde and staining with Giemsa, invaded cells on the bottom surface were counted in six randomly chosen fields at a magnification of 200×. For every condition, tests were conducted in triplicate.

### Statistical analysis

All results are presented as mean values ± standard deviation (SD), based on a minimum of three independent experimental replicates. Statistical analyses were performed using GraphPad Prism software (version 9.4.0; GraphPad Software). Statistical differences between paired groups were analyzed using a two-tailed Student’s t-test, whereas one-way or two-way ANOVA followed by suitable post hoc tests was employed for multi-group comparisons. Statistical significance was defined as p < 0.05.

## Results

### Efficient silencing of SOX2OT is associated with increased miR-143 expression in NSCLC cells

Cocktail siRNAs targeting human SOX2OT were used to suppress SOX2OT expression in Calu-3 and A549 cell lines to investigate the biological function of SOX2OT in NSCLC. Fluorescence microscopy utilizing dye-labeled control siRNA verified transfection effectiveness, and qRT-PCR analysis showed a significant decrease in SOX2OT expression in comparison to PBS-treated and scrambled control cells (Figure 1A). We next assessed the effect of SOX2OT depletion on miR-143 expression. SOX2OT knockdown resulted in a significant upregulation of miR-143 in both NSCLC cell lines, whereas no change was observed in control groups. The results reveal a negative correlation between SOX2OT and miR-143 expression in NSCLC cells (Figure 1B). Using the IntaRNA tool (version 3.4.1) (20), a putative direct interaction between SOX2OT and miR-143 was predicted. The analysis revealed complementary binding regions capable of forming multiple base-pair interactions, with a minimum free energy of −11.82 kcal/mol and a hybridization energy of −23.17 kcal/mol, supporting the thermodynamic stability of the predicted SOX2OT–miR-143 interaction (Figure 1C).

**Figure 1.**
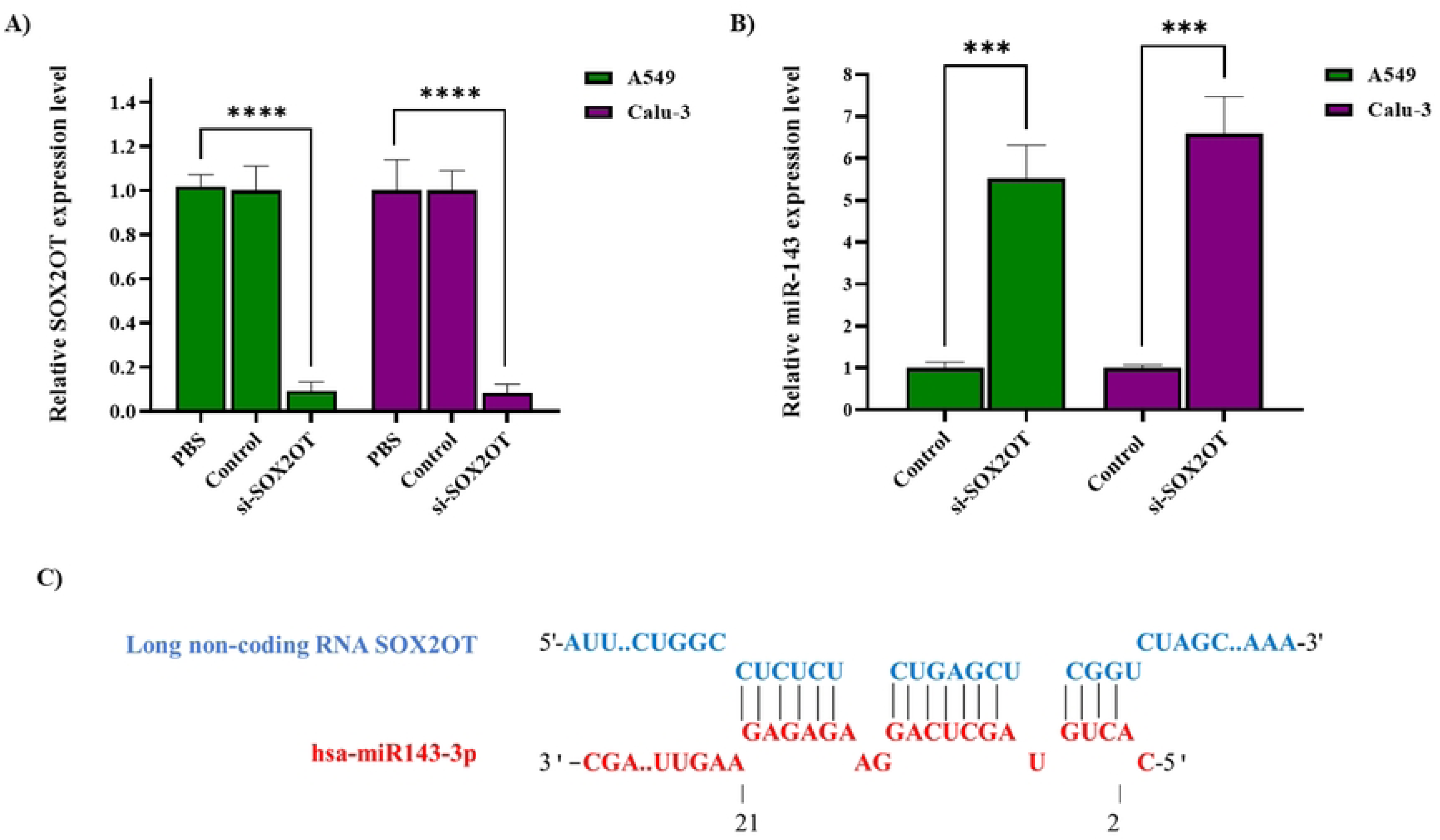
Knockdown of SOX2OT increases miR-143 expression in NSCLC cell lines. **A)** Confirmation of SOX2OT silencing efficiency in Calu-3 and A549 cells. qRT-PCR analysis of SOX2OT expression levels 48 h post-transfection with SOX2OT-targeting siRNA cocktail compared to untreated cells and scrambled siRNA control (n = 3 independent experiments; mean ± SD; ****p < 0.0001). **B)** Effect of SOX2OT knockdown on miR-143 expression. qRT-PCR analysis of miR-143 levels in Calu-3 and A549 cells 48 h following transfection with si-SOX2OT or the NC control (n = 3 independent experiments; mean ± SD; ***p < 0.001). **C)** In silico prediction of direct interaction between SOX2OT and miR-143 using IntaRNA (v3.4.1). Schematic representation of the predicted binding sites with complementary base-pairing, minimum free energy (MFE = −11.82 kcal/mol), and hybridization energy (−23.17 kcal/mol).

### SOX2OT depletion modulates miR-143–associated oncogenic signaling pathways

To investigate the downstream molecular effects of SOX2OT depletion, we analyzed the expression of in silico–predicted miR-143 target genes that are implicated in NSCLC pathobiology. Quantitative real-time PCR indicated that silencing of SOX2OT resulted in a notable reduction in the transcriptional abundance of STAT3, EZH2, and CXCL13 in both Calu-3 and A549 cells. Conversely, PTEN expression was markedly upregulated following SOX2OT knockdown (Fig. 2A, B). Western blotting further validated the mRNA findings, revealing a pronounced decrease in STAT3, and CXCL13 protein levels following SOX2OT silencing. Consistent protein loading was confirmed by the constant expression of β-actin in all groups (Fig. 2C). All of these findings suggest that SOX2OT influences important oncogenic signaling elements in NSCLC cells, possibly via a regulatory axis connected to miR-143.

**Figure 2.**
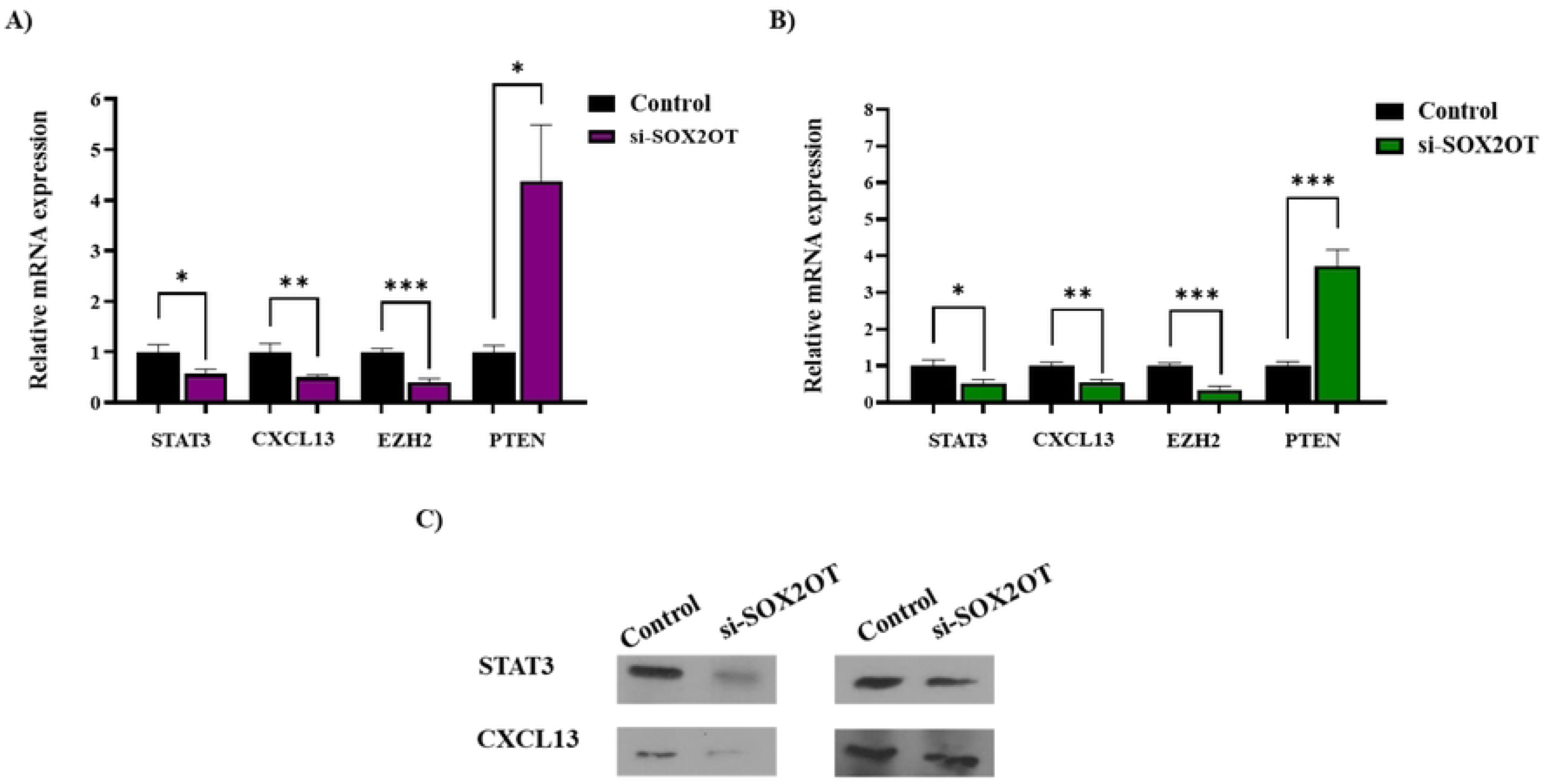
In NSCLC cells, SOX2OT knockdown controls oncogenic signaling associated with miR-143. **A, B)** qRT-PCR study demonstrating the expression levels of STAT3, EZH2, CXCL13, and PTEN mRNA expression in A549 (A) and Calu-3 (B) cells treated with either si-SOX2OT or NC. STAT3, EZH2, and CXCL13 transcripts were significantly downregulated when SOX2OT was silenced, but PTEN expression was noticeably increased in both NSCLC cell lines. **C)** Representative Western blot images showing STAT3, and CXCL13 protein expression in A549 (left) and Calu-3 (right) cells following SOX2OT silencing. β-actin was used as the internal loading control to guarantee consistent protein loading among samples. Results are expressed as mean ± SD derived from a minimum of three separate experiments. Differences between groups were evaluated with a two-tailed Student’s t-test (*p < 0.05, **p < 0.01, *p < 0.001).

### SOX2OT promotes NSCLC cell proliferation and regulates G1-phase cell cycle progression

48 hours after siRNA transfection, flow cytometry was utilized to assess if SOX2OT influences cell cycle progression. G1-phase arrest was induced in both NSCLC cell lines by SOX2OT knockdown, which resulted in a notable concentration of cells in the G1 phase together with a reduction in S and G2/M phase populations (Fig. 3A, B). The trypan blue exclusion experiment verified that SOX2OT knockdown dramatically reduced proliferation, as seen by a notable decrease in viable cell counts, which is consistent with these cell cycle changes (Fig. 3C). All of these results point to SOX2OT’s positive regulation of NSCLC cell cycle progression and promotion of cellular proliferation.

**Figure 3.**
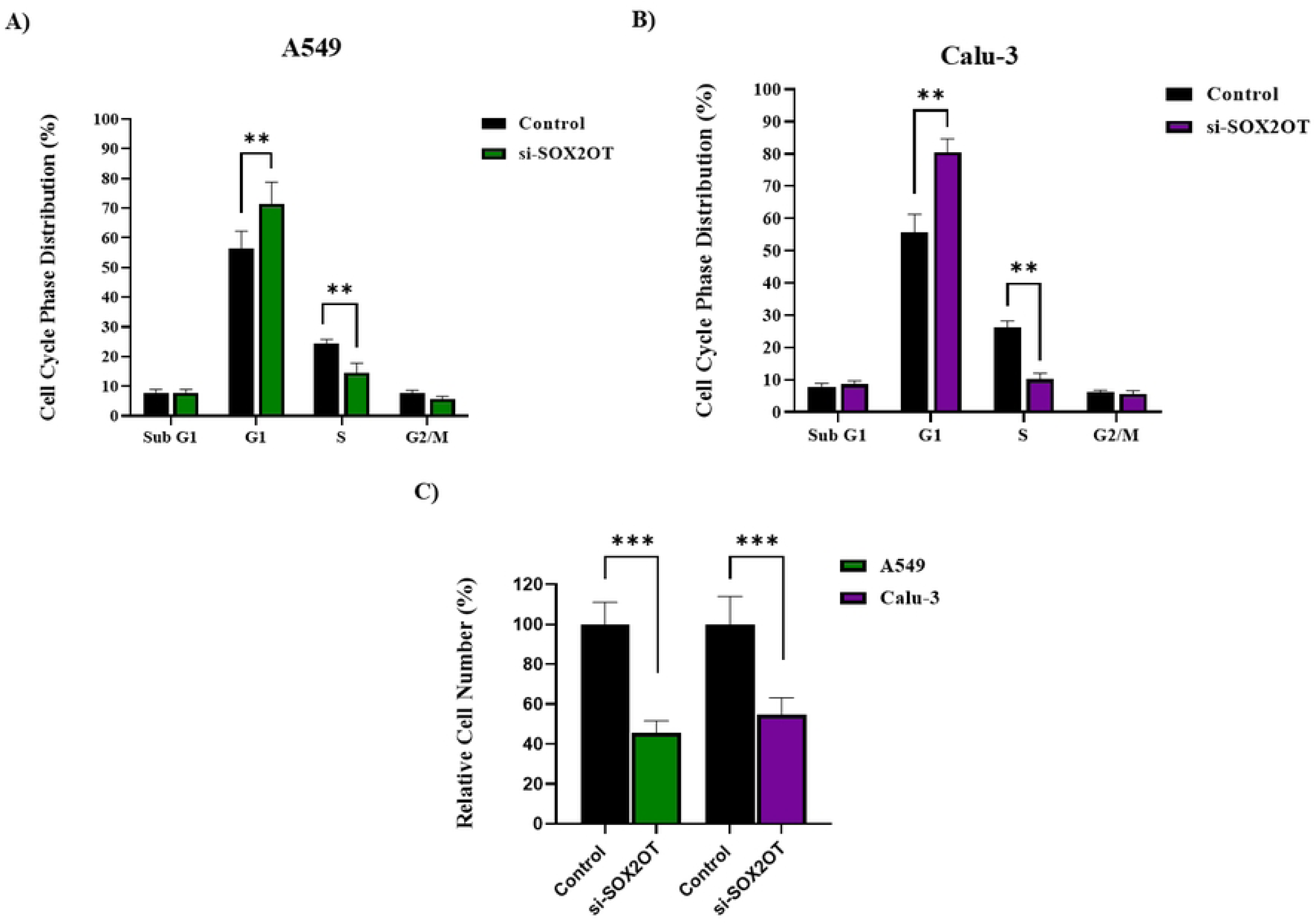
SOX2OT increases NSCLC cells’ ability to proliferate and controls the course of the cell cycle. **A, B)** 48 hours after transfection with si-SOX2OT or negative control, cell cycle progression was studied by flow cytometry distribution in Calu-3 and A549 cells. G1-phase arrest was shown by a merked rise in the percentage of cells in the G1 phase following SOX2OT silencing, which was followed by a similar drop in S-phase cells. C) Cell proliferation was evaluated using the trypan blue assay, demonstrating that the depletion of SOX2OT markedly reduced the number of viable cells. Furthermore, 48 hours after si-SOX2OT transfection, the MTT assay findings further demonstrated reduced proliferative activity in both A549 and Calu-3 cells. Results are expressed as mean ± SD obtained from a minimum of three separate experiments. Statistical significance was considered at the levels of *p < 0.01 and p < 0.001.

### SOX2OT knockdown enhances apoptotic cell death in NSCLC cells

AO/EtBr staining was initially utilized to evaluate the effect of SOX2OT knockdown on apoptotic cell death. Relative to control groups, fluorescence microscopic examination revealed a pronounced elevation in apoptotic cells in SOX2OT-depleted NSCLC cells, distinguished by hallmark apoptotic morphologies (Fig. 4A). Quantitative confirmation was obtained through Annexin V–FITC/PI dual labeling and flow cytometry. In comparison to control cells, SOX2OT depletion caused a significant increase in the total apoptotic fraction at 48 hours after transfection, encompassing both early and late phases of apoptosis (Fig. 4B, C). All of these findings point to a possible function for SOX2OT in providing resistance to apoptosis in NSCLC cells, as reduction of SOX2OT increases apoptotic cell death.

**Figure 4.**
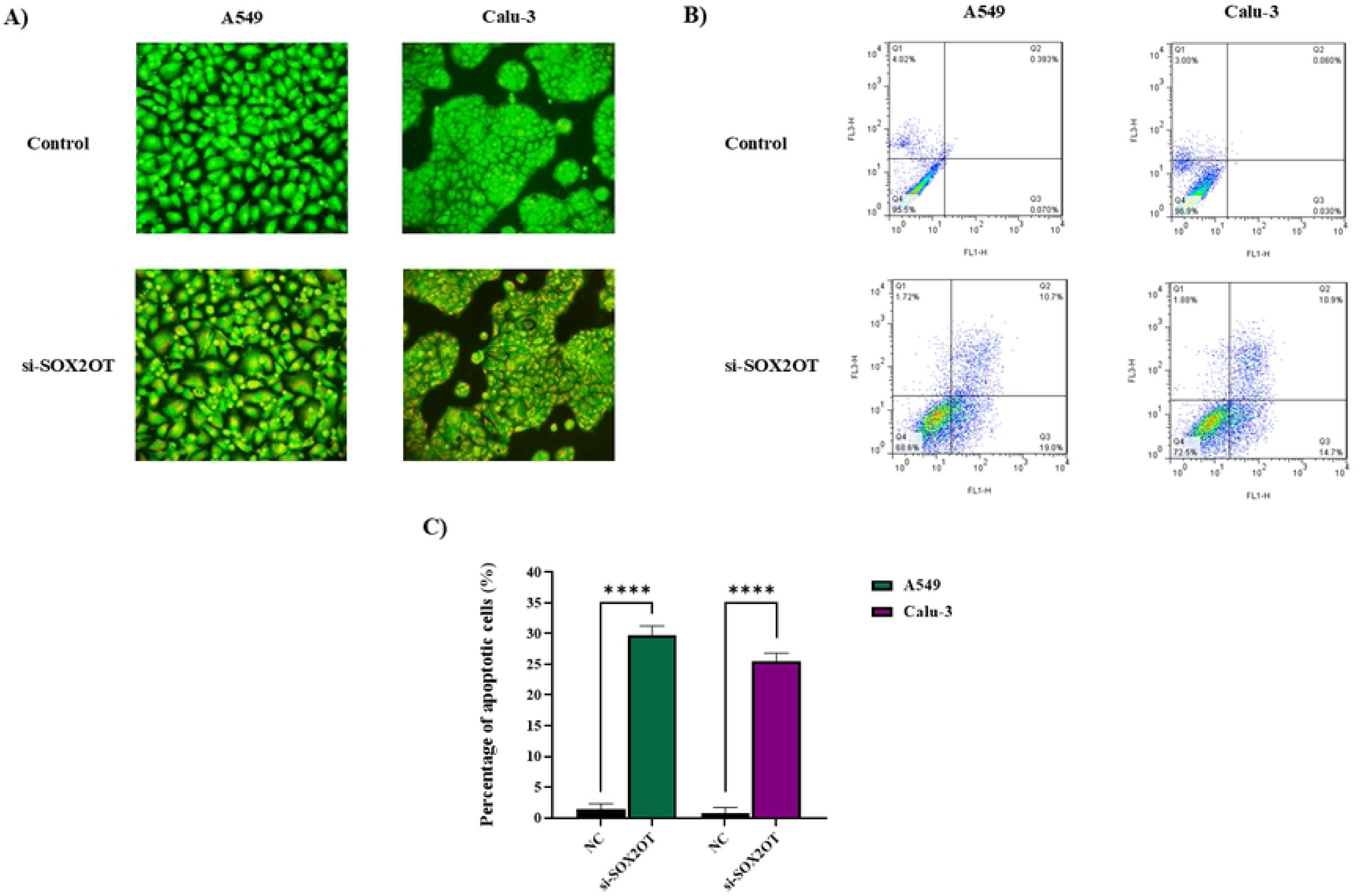
SOX2OT knockdown promotes apoptotic cell death in NSCLC cells. **A)** 48 hours post-introduction of si-SOX2OT or a scrambled control, Calu-3 and A549 cells underwent staining with AO/EtBr, showing a higher percentage of apoptotic cells after SOX2OT silencing. **B)** FlowJo software was used to examine representative flow cytometry plots of Annexin V-FITC/PI-stained cells that show apoptotic cell populations in the control and SOX2OT-depleted groups. **C)** Quantitative analysis of early and late apoptotic fractions at 48 hours after knockdown, showing a significant increase in apoptosis upon SOX2OT suppression. Results are expressed as mean ± SD derived from at least three separate experiments. Differences were regarded as statistically significant at *p < 0.05 and **p < 0.01 compared with the control group.

### SOX2OT silencing suppresses NSCLC cell migration

To determine whether SOX2OT contributes to NSCLC cell motility, wound-healing analysis was performed. SOX2OT-silenced A549 and Calu-3 cells exhibited significantly slower wound closure than control cells, supporting a marked decrease in migratory activity after SOX2OT suppression (Fig. 5A-C). A transwell migration experiment was used to confirm these results. In both NSCLC cell lines, SOX2OT knockdown dramatically decreased the amount of migratory cells in comparison to control groups, which is consistent with the wound healing results (Fig. 5D, E). All of these findings show that SOX2OT stimulates NSCLC cell motility and that its knockdown successfully inhibits migratory activity in vitro.

**Figure 5.**
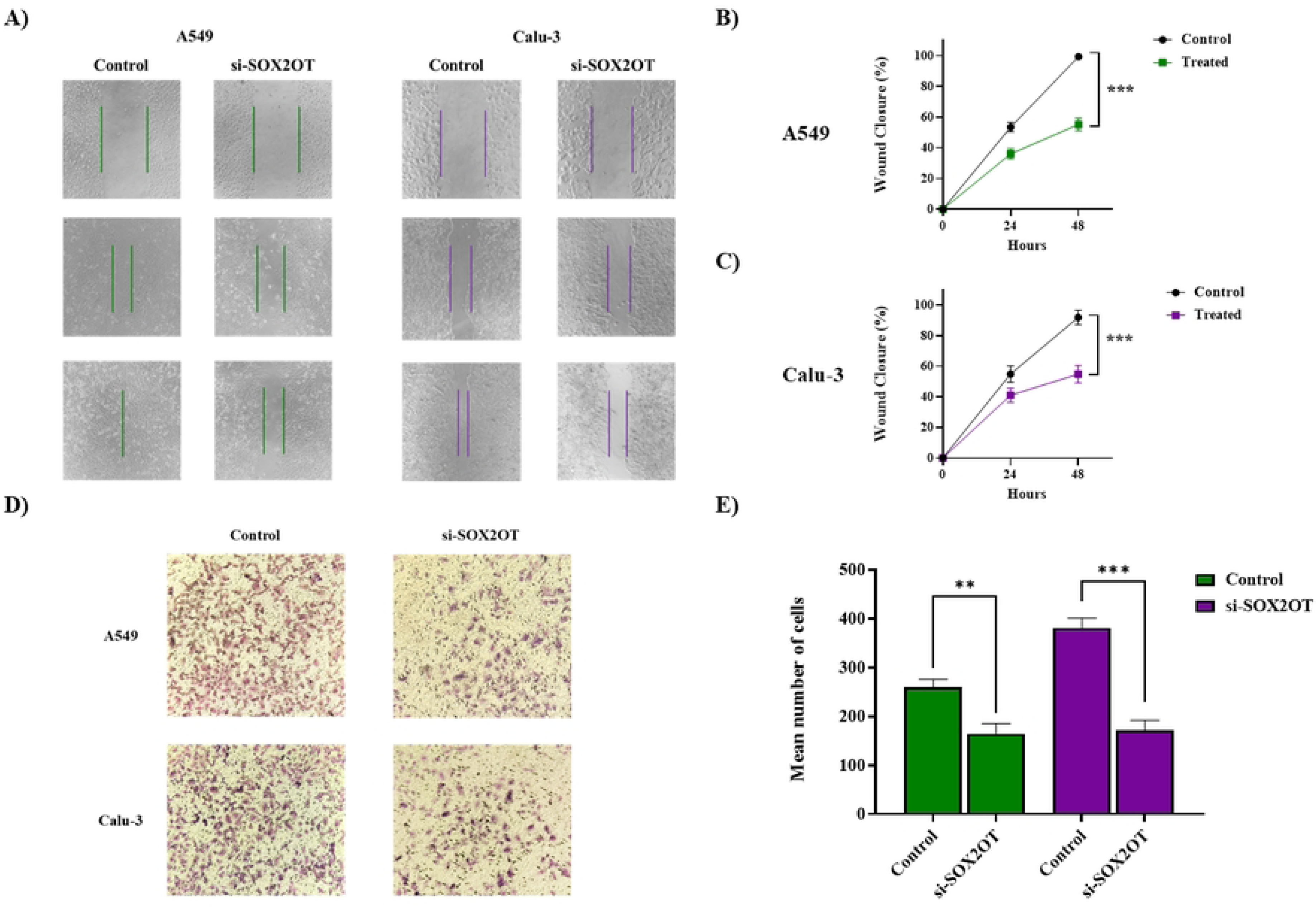
SOX2OT silencing suppresses migratory capacity of NSCLC cells. **A, B)** Scratch wound healing assay representative micrographs and quantitative analysis showing decreased wound closure in Calu-3 and A549 cells after si-SOX2OT transfection in comparison to scrambled siRNA controls. Pictures were taken at 0, 24, and 48 hours following the creation of the wound. **C)** The transwell migration experiment demonstrates that SOX2OT-depleted A549 and Calu-3 cells move significantly fewer cells than control groups. Quantification of migrating cells is accompanied by representative photos. The mean ± SD of at least three separate experimental repetitions is used to express the data. SOX2OT controls EMT in NSCLC cells and inhibits NSCLC cell invasion.

SOX2OT knockdown significantly decreased the number of invasive cells in Matrigel-coated transwell invasion experiments (Fig. 6A, B), indicating that SOX2OT promotes both migration and invasion in NSCLC cells. We looked at how SOX2OT regulates the EMT, which is crucial for the spread of tumors. SOX2OT knockdown dramatically reduced the EMT-inducing TFs SNAIL and TWIST in both cell lines, according to qRT-PCR analysis. On the other hand, this silencing caused the mesenchymal markers N-cadherin and vimentin to be downregulated and the epithelial marker E-cadherin to be upregulated (Fig. 6C, D).

**Figure 6.**
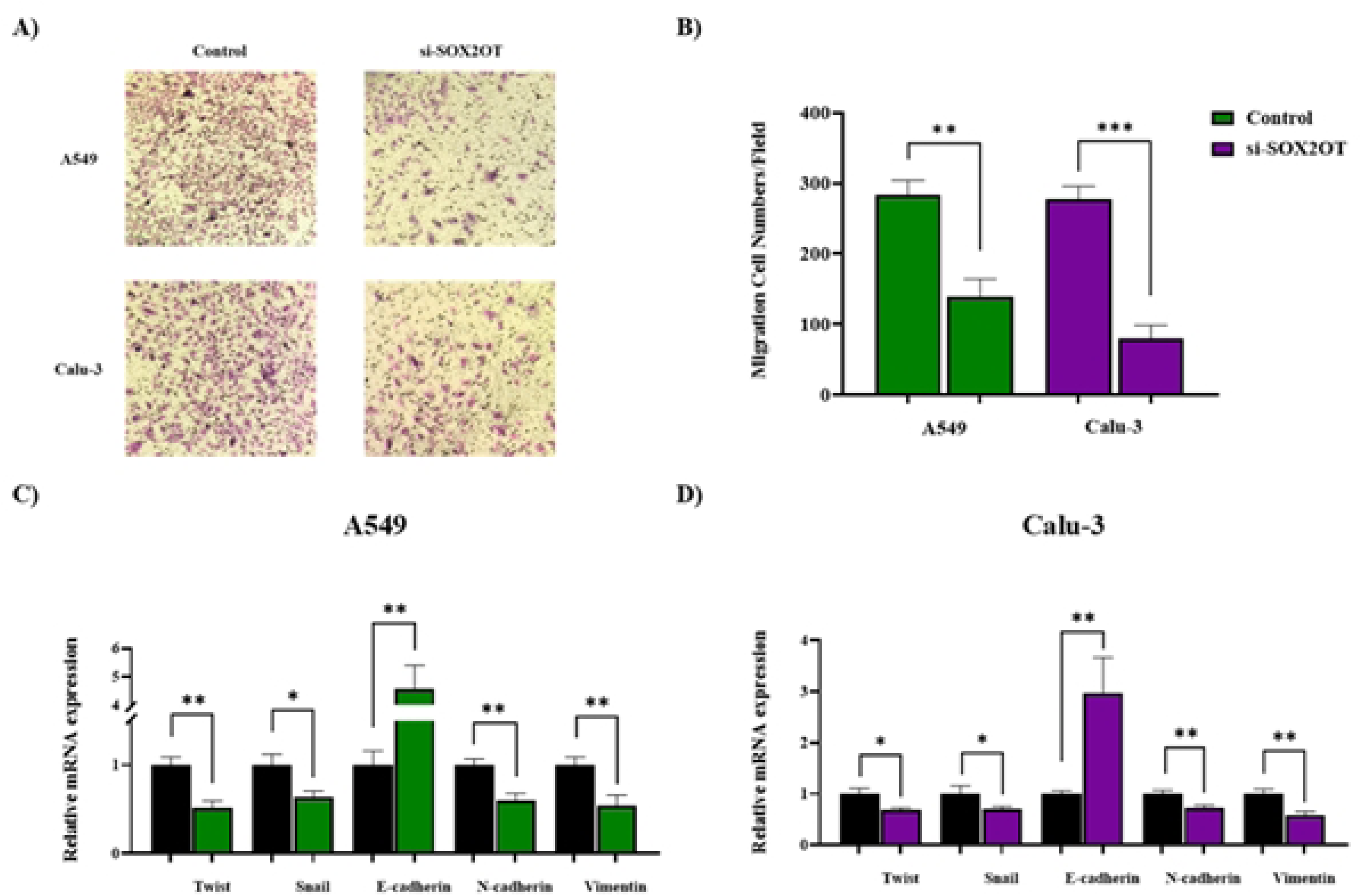
SOX2OT enhances invasive behavior and modulates EMT in NSCLC cells. **A)** Representative micrographs from Matrigel-coated transwell invasion assays illustrating diminished invasive potential in Calu-3 and A549 cells following SOX2OT knockdown compared with scrambled siRNA controls. **B)** Quantitative evaluation of invaded cells confirming a significant reduction in invasive capacity upon SOX2OT silencing in both NSCLC cell lines. **C)** After treatment with si-SOX2OT and following reduction of SOX2OT, qRT-PCR analysis of EMT-related gene expression in A549 cells demonstrated diminished levels of the TFs SNAIL and TWIST, elevated expression of the epithelial marker E-cadherin, and reduced expression of the mesenchymal markers Vimentin and N-cadherin. D) Calu-3 cells showed similar EMT-related expression patterns, with downregulation of SNAIL and TWIST, elevation of E-cadherin, and suppression of N-cadherin and Vimentin following SOX2OT silencing. The information is displayed as mean ± SD. *p < 0.05, **p < 0.01, and ***p < 0.001 denote statistical significance.

## Discussion

NSCLC remains a leading cause of cancer-related mortality, largely due to aggressive tumor behavior, early metastasis, and limited durable responses to current therapies (21, 22). Increasing evidence indicates that lncRNAs play central roles in coordinating oncogenic signaling networks that drive these malignant traits (8). According to this study, SOX2OT is a crucial regulator of the development of NSCLC. Our results imply that the modification of a miR-143-dependent regulatory axis is at least partially responsible for the carcinogenic effects of SOX2OT. In addition to inducing G1-phase arrest and death, silencing SOX2OT has been demonstrated to significantly reduce proliferation, migration, invasion, and EMT in NSCLC cells. This characteristic is in line with previous results that identified SOX2OT as an oncogenic lncRNA in lung cancer and other malignancies, where it contributes to cell survival, metastatic potential, and tumor growth (7, 9). Importantly, our study expands the current understanding of SOX2OT function in NSCLC by linking it to miR-143, a well-established tumor-suppressive microRNA.

It has been well documented that MiR-143 is downregulated in NSCLC and that it targets several oncogenic pathways to impede cell proliferation, migration, invasion, and EMT (15, 16). In line with these observations, we found that SOX2OT depletion leads to a robust upregulation of miR-143 in two independent NSCLC cell lines. This inverse relationship suggests that SOX2OT may negatively regulate miR-143 availability, potentially through a competing endogenous RNA–like mechanism, as has been described for SOX2OT in other cancer contexts. Although direct molecular interaction assays are required to definitively establish this mechanism, our data suggests a functional linkage between SOX2OT and miR-143 in NSCLC cells.

At the molecular level, we identified several key oncogenic effectors downstream of the SOX2OT/miR-143 axis, including STAT3, EZH2, and CXCL13, alongside restoration of the tumor suppressor PTEN. STAT3 is a key TF, which promotes tumor cell proliferation, survival, immune modulation, and EMT, and its aberrant activation is a hallmark of aggressive NSCLC. The observed downregulation of STAT3 following SOX2OT silencing provides a plausible mechanistic basis for the reduced proliferative capacity and increased apoptotic susceptibility of NSCLC cells in this study.

Tumor suppressor genes are epigenetically silenced by EZH2, a catalytic component of the Polycomb Repressive Complex 2, which is often overexpressed in non-small cell lung cancer (23, 24). Our data indicate that SOX2OT depletion suppresses EZH2 expression and is accompanied by increased PTEN levels, suggesting relief of EZH2-mediated repression. Given the central role of the PTEN/PI3K/AKT axis in regulating cell survival and apoptosis (25), this regulatory cascade likely contributes to the pronounced apoptotic response observed upon SOX2OT knockdown.

CXCL13 has emerged as an important chemokine in cancer biology, influencing tumor cell migration, invasion, and interactions with the tumor microenvironment (26). Although its role in NSCLC is less extensively characterized than STAT3 or EZH2, accumulating evidence implicates CXCL13 signaling in promoting metastatic behavior and immune modulation (27). The downregulation of CXCL13 following SOX2OT silencing suggests that SOX2OT may also influence chemokine-driven pathways that support NSCLC progression (28).

At the phenotypic level, our findings demonstrate that SOX2OT is a strong positive regulator of EMT in NSCLC cells. SOX2OT knockdown suppressed EMT-inducing TFs such as SNAIL and TWIST, restored epithelial marker expression, and reduced mesenchymal marker levels. These molecular changes were accompanied by significant suppression of cell migration and invasion, underscoring the functional relevance of SOX2OT-driven EMT programming. Given the pivotal role of EMT in metastatic dissemination and therapeutic resistance, these observations position SOX2OT as an important upstream coordinator of metastatic potential in NSCLC.

Despite the strength of the in vitro evidence presented here, several limitations should be acknowledged. Although previous studies has demonstrated direct binding between SOX2OT and miR-143 using luciferase reporter assays in a different biological context (19, 29), direct mechanistic validation was not conducted in the present study. Therefore, while our bioinformatics prediction, expression correlation, and functional data strongly support a regulatory relationship in NSCLC, the precise ceRNA mechanism in lung cancer cells requires further confirmation. Second, all functional assays were conducted in only two NSCLC cell lines (Calu-3 and A549 and), which may limit the broader applicability of the findings across diverse NSCLC subtypes and genetic backgrounds. Third, the present study is restricted to in vitro experimental systems and does not include in vivo validation using xenograft or orthotopic tumor models. Such models would be essential to further substantiate the role of the SOX2OT/miR-143 regulatory axis in tumor growth and metastatic progression within a more physiologically relevant microenvironment. Fourth, the clinical relevance remains preliminary because SOX2OT and miR-143 expression levels were not analyzed in a large cohort of NSCLC patient samples, nor correlated with clinicopathological features or patient survival outcomes. Finally, potential off-target effects of siRNA-mediated knockdown, although minimized by using a cocktail of siRNAs, cannot be completely ruled out. Future studies incorporating NSCLC-specific binding validation, in vivo models, and comprehensive clinical correlation analyses will be crucial to successful establish the therapeutic potential of targeting the SOX2OT/miR-143 axis in NSCLC.

In summary, this study identifies SOX2OT as a central oncogenic lncRNA that promotes NSCLC progression by modulating a miR-143–associated regulatory network involving STAT3, EZH2, CXCL13, and PTEN. By integrating post-transcriptional and epigenetic regulation, SOX2OT coordinates multiple hallmarks of malignancy, including uncontrolled proliferation, apoptosis resistance, EMT, and invasive behavior. These results demonstrate SOX2OT as a potentially useful molecular vulnerability in NSCLC and offer compelling evidence for additional research into SOX2OT-targeted lung cancer treatment approaches (Fig. 7).

**Figure 7.**
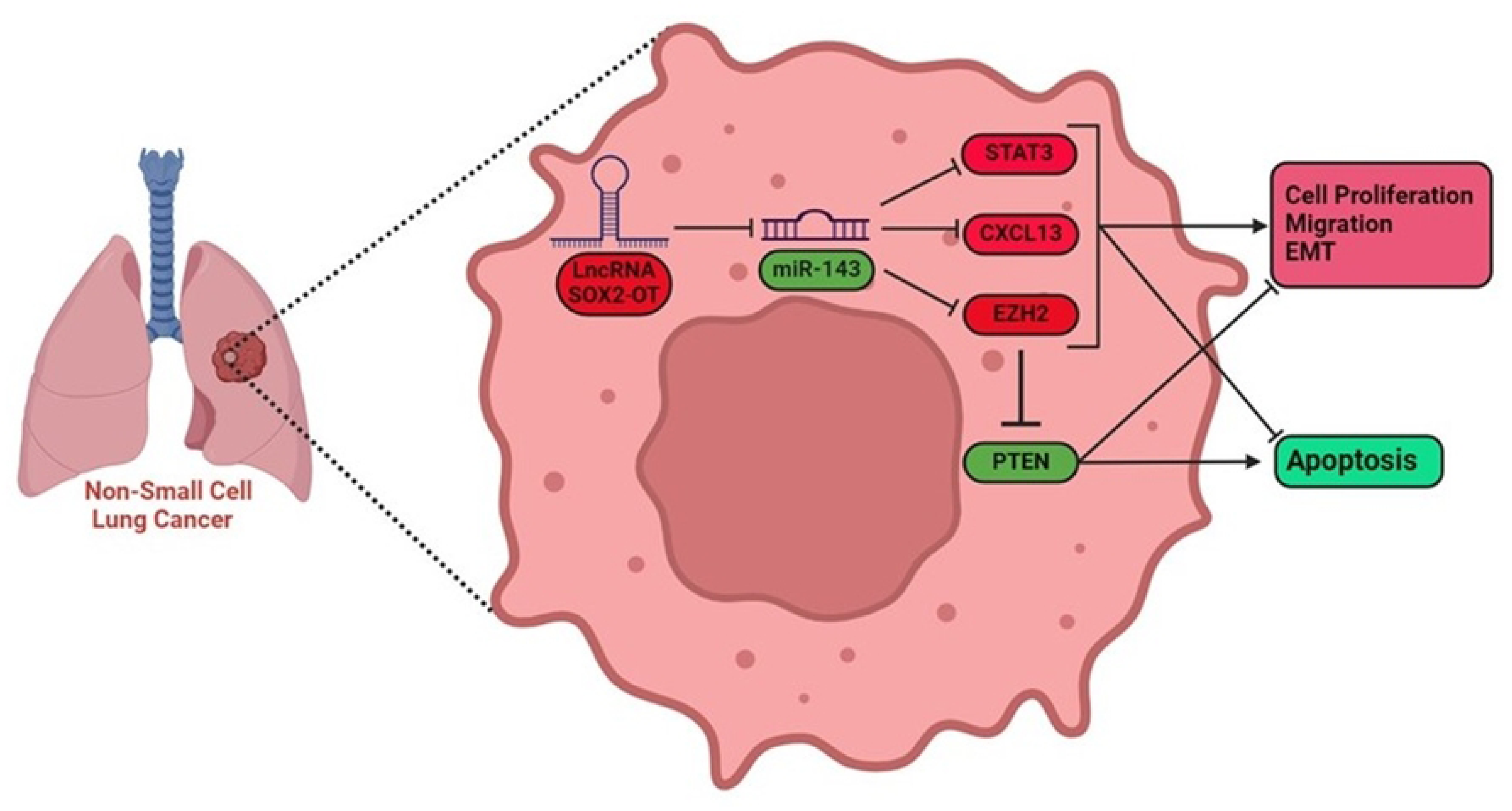
Proposed model of SOX2OT-mediated regulation of NSCLC progression. Schematic representation of the proposed regulatory network through which the long non-coding RNA SOX2OT contributes to NSCLC progression. Elevated SOX2OT expression is associated with reduced availability of miR-143-3p, potentially through a competing endogenous RNA–like mechanism. Decreased miR-143-3p levels are accompanied by increased expression of oncogenic effectors, including STAT3, EZH2, and CXCL13, and reduced expression of the tumor suppressor PTEN. Activation of these pathways facilitates malignant progression in NSCLC by enhancing proliferation, EMT, migration, and invasion, while suppressing apoptosis. Dashed lines indicate potential or indirect regulatory links based on combined *in vitro* and *in silico* analyses.

## Conclusion

The lncRNA SOX2OT is identified in this work as a key regulator of malignant phenotypes in non-small cell lung cancer. We demonstrate that SOX2OT inhibits apoptotic cell death while simultaneously promoting NSCLC cell proliferation, survival, migration, invasion, and EMT through integrated molecular and functional studies. Mechanistically, these tumor-promoting effects seem to operate through a miR-143–orchestrated signaling cascade that regulates critical downstream targets, including STAT3, EZH2, CXCL13, and the tumor suppressor PTEN. Our findings expand the current understanding of lncRNA-mediated post-transcriptional and epigenetic regulation in NSCLC and highlight SOX2OT as an important upstream coordinator of signaling pathways that govern tumor growth and metastatic potential. By integrating miRNA regulation with oncogenic transcriptional and epigenetic programs, SOX2OT emerges as a potential molecular vulnerability in NSCLC. Collectively, this work provides a strong rationale for further investigation of the SOX2OT/miR-143 axis as a therapeutic target in lung cancer. Future studies incorporating direct molecular interaction assays, in vivo validation, and clinical correlation analyses will be essential to establish the translational relevance of SOX2OT-targeted strategies and to determine their potential utility in improving NSCLC patient outcomes.

## Declarations

### Ethics approval and consent to participate

Not applicable (established cell lines only).

### Author Contributions

**Mohadeseh Zarei**: Conceptualization, Methodology, Investigation, Formal analysis, Visualization, Writing – original draft, Writing – review & editing. **Elahe Asadollahi**: Investigation, Methodology, Validation, Visualization, Data curation, Writing – review & editing. **Babak Jahangiri**: Investigation, Methodology, Formal analysis, Visualization. **Jamshid Raheb**: Conceptualization, Supervision, Project administration, Funding acquisition, Writing – review & editing. All authors read and approved the final manuscript.

